# Efficient Through-Bond Propagation of Nuclear-Spin Hyperpolarization via TOCSY-Enhanced LC-Photo-CIDNP

**DOI:** 10.64898/2026.09.11.751018

**Authors:** Ji Ho Jeong, Anubhab Halder, Silvia Cavagnero

## Abstract

Nuclear magnetic resonance (NMR) spectroscopy provides atomic-resolution insights into biomolecular structure, dynamics and interactions under non-perturbative conditions. However, the widespread applicability of this technique continues to be limited by its intrinsically low sensitivity. This drawback is particularly severe in the case of concentration-limited samples. Low-concentration photochemically induced dynamic nuclear polarization (LC-photo-CIDNP) is an optically enhanced NMR hyperpolarization strategy capable of boosting NMR sensitivity *in situ* by several orders of magnitude under physiologically relevant environments. Yet, the inherent nature of the LC-photo-CIDNP phenomenon confines the achievable enhancements to only a subset of nuclei, thus limiting the extent of attainable residue-specific information. Here, we overcome this shortcoming by integrating LC-photo-CIDNP with Total Correlation Spectroscopy (TOCSY) to propagate initial hyperpolarization to other nuclei via scalar-coupled spin networks. First, we demonstrate efficient through-bond transfer of LC-photo-CIDNP enhanced signals throughout entire intramolecular atomic frameworks via 1D ¹H- and ¹³C-detected experiments. Second, we extend the approach to a model protein and demonstrate efficient polarization transfer via 1D and 2D ¹H-detected ¹³C-TOCSY LC-photo-CIDNP. Intrinsically photo-CIDNP-inactive nuclei can thus be visualized via ^13^C-^1^H correlations within minutes down to 10 μM concentration. In all, this work establishes a general strategy to achieve through-bond propagation of LC-photo-CIDNP hyperpolarization. This technology enables gaining residue-specific structural insights on amino acids and proteins at previously unattainable low concentrations.

**Significance Statement:** Nuclear magnetic resonance (NMR) plays a key role in the elucidation of biomolecular structure, dynamics and interactions. Yet, it is extremely insensitive. Here, we synergistically combine an optically enhanced NMR technology known as LC-photo-CIDNP with total correlation spectroscopy (TOCSY). Via this approach, we achieve highly enhanced detection thresholds, together with dramatic increases in the number of hyperpolarized nuclei. This technology is readily applicable to both amino acids and proteins in solution. Remarkably, TOCSY-enhanced LC-photo-CIDNP enables a tunable degree of through-bond hyperpolarization propagation, with no restrictions in biomolecular tumbling or compaction. In all, this technology enables unprecedented NMR sensitivity while encompassing a broad set of nuclei, thus highlighting versatile routes towards a more comprehensive structural characterization of biomolecules at low concentration.

## Introduction

Solution State Nuclear magnetic resonance (NMR) spectroscopy is versatile tool to elucidate biomolecular structure, dynamics, interactions and allosteric transitions at atomic resolution under physiologically relevant conditions. Despite decades of technological advances, including magnets operating at GHz frequencies and cryogenic probes, poor signal-to-noise remains a persistent, critical bottleneck.(1–3) The low sensitivity of NMR arises from the inherently small nuclear spin polarization at thermal equilibrium.(4, 5) This challenge is particularly pronounced in the case of inherently low-abundance molecules or systems that need to be kept dilute due to high aggregation propensities.(6) The small energy differences between the nuclear spin states at equilibrium in the presence of a magnetic field is a fundamental constraint of NMR.(4, 6) This limitation prompted the development of a variety of spin hyperpolarization approaches aimed at generating transient non-Boltzmann nuclear-spin populations (Fig. 1A).(4, 7–10)

**Figure 1.**
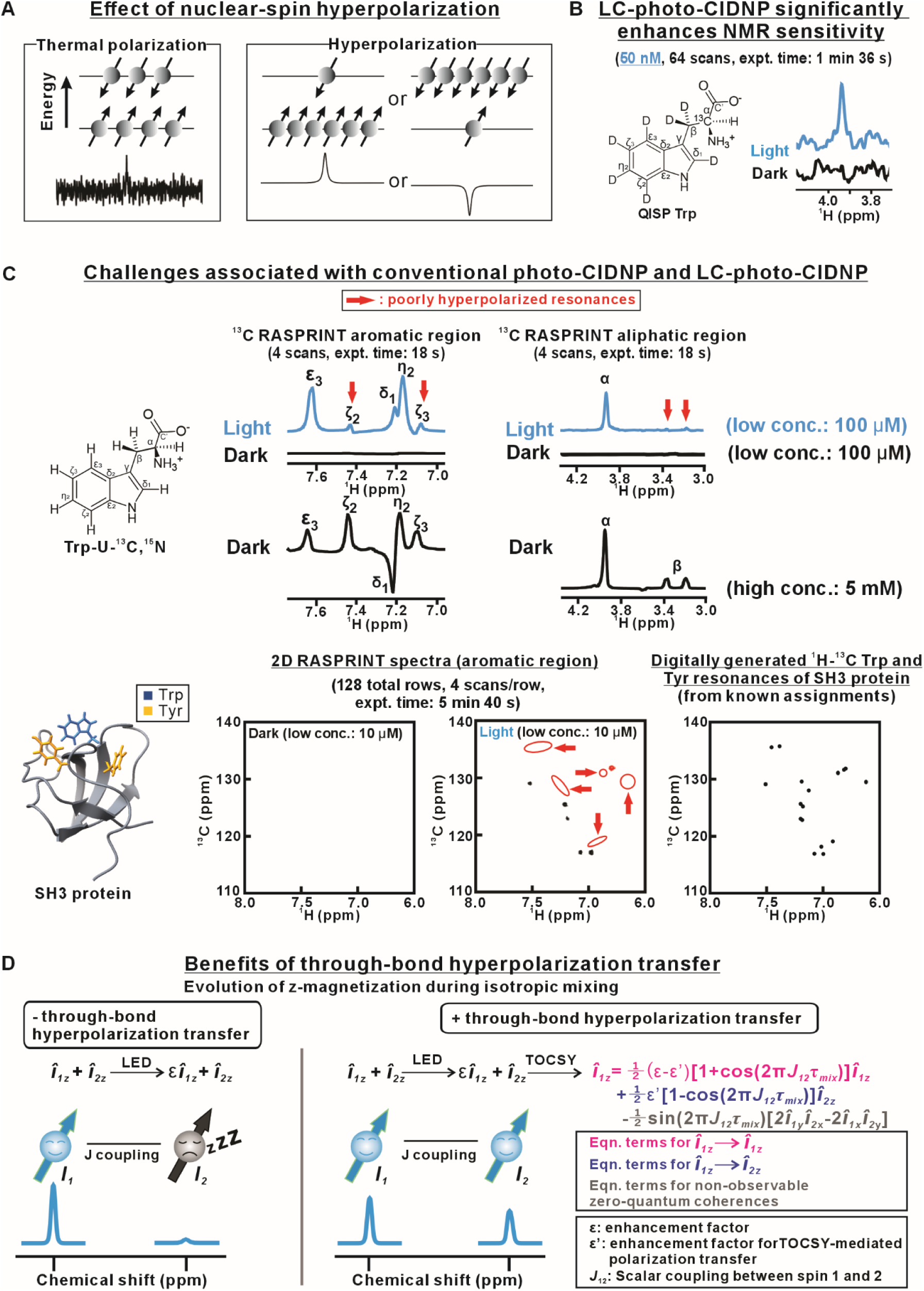
Some limitations of LC-photo-CIDNP hyperpolarization can be overcome via TOCSY-mediated hyperpolarization transfer. *(A)* Schematic illustration highlighting the basic concept underlying nuclear-spin hyperpolarization. This process generates a large population imbalance between nuclear spin states, resulting in dramatically enhanced positive or negative NMR resonances. *(B)* LC-photo-CIDNP enables highly sensitive NMR detection. Representative ^1^H detected ^13^C LC-photo-CIDNP spectrum of QISP Trp acquired under light conditions. The data demonstrate substantial signal enhancements at nM sample concentration. *(C)* Limitations of conventional LC-photo-CIDNP. Only a subset of Trp-U-^13^C,^15^N resonances experience hyperpolarization, while several others remain weakly enhanced or altogether undetectable (red arrows), under light conditions. The same concept applies to the LC-photo-CIDNP RASPRINT spectrum of the SH3 protein under light conditions. Several expected resonances are absent, as assessed upon comparison with the digitally generated ^1^H–^13^C Trp and Tyr resonances based on tknown assignments.(53) *(D)* Basic concepts underlying TOCSY-mediated through-bond hyperpolarization transfer. The equations describe hyperpolarization transfer between two J-coupled nuclear spins. All LC-photo-CIDNP experiments were carried out in 10 mM potassium phosphate (pH ∼7.2). The QISP Trp, Trp-U-¹³C, ¹⁵N, and SH3 samples contained 2.5 μM, 25 μM, and 10 μM fluorescein, respectively. See Materials and Methods for details.

Among the presently available hyperpolarization tools, photochemically induced dynamic nuclear polarization (photo-CIDNP) has the unique ability to rapidly generate (∼0.2-1 s) hyperpolarization of solvent-exposed target molecules that are susceptible to reversible electron transfer. The latter include free aromatic amino acids either in isolation or within proteins. Two of the major strengths of the photo-CIDNP technology are its cost-effective nature and the fact that it can be carried out *in situ* under physiologically relevant conditions.(11–16) At its core, liquid-state photo-CIDNP relies on the productive diffusional encounters between the molecule of interest and the photoexcited triplet state of a dye. These collisions are followed by rapid (ca. ps timescale) electron transfer, leading to the formation of transient triplet-state radical pairs within a solvent cage.(17–20) The latter undergo nuclear-spin-state-dependent triplet-singlet (TS) mixing. This process lies at the origin of the ultimately observed large sensitivity enhancements. The TS mixing proceeds at a frequency, corresponding to the difference in the electron paramagnetic resonance (EPR) frequencies of the two unpaired electrons within the radical pair.(21, 22) A nucleus that enjoys an appreciable hyperfine coupling to the unpaired electron modulates the EPR precession frequency of the unpaired electron delocalized across the pertinent component of the radical pair. Importantly, the extent of this modulation depends on whether that nucleus (e.g., consider a spin ^1^⁄_2_ nucleus, e.g., ^1^H or ^13^C) is in an α (corresponding to a nuclear spin quantum number, *m_I_* = + ^1^⁄_2_) or β (*m_I_* = − ^1^⁄_2_) spin state. As a result of the different EPR precession frequencies of the α and β states, a non-Boltzmann distribution of nuclear spin states is ultimately generated within the recombination products.(4, 22, 23) This hyperpolarization manifests itself as strongly enhanced absorptive or emissive NMR signals.(23, 24)

In recent years, a low-concentration version of photo-CIDNP (a.k.a. LC-photo-CINDP) has been developed through a series of methodological advances that augment the sensitivity of the technique, providing access to previously unattainable low concentrations.(14, 16, 25–28) The key elements of LC-photo-CIDNP include: (i) use of photosensitizer dyes with a long photoexcited triplet-state lifetime,(17) (ii) inclusion of an enzyme system for the facile depletion of molecular oxygen,(29) (iii) use of reductive radical quenchers to enable prolonged data collection on dilute (≤ low μM) samples,(28) (iv) development of homo and heteronuclear hyperpolarization and signal-detection schemes optimized for ultrafast data collection,(7, 11, 30) and (v) implementation of selective isotope-labeling to further enhance hyperpolarization.(14, 26, 27, 31) Together, these innovations extended the detection limits of photo-CIDNP down to the 20 nM regime within minutes.(26, 32) A representative example, showing the spectrum of the Trp amino acid, which bears a quasi-isolated spin pair (QISP Trp), is shown in Fig. 1B. LC-photo-CIDNP is particularly advantageous in biomolecular NMR, given that many intra- and extra-cellular molecules are often available at low-μM-to-pM levels under physiologically relevant conditions. In all, the capability to probe biomolecular processes at low concentration facilitates the *in situ* characterization of sparsely populated conformational states, transient intermediates, and protein–ligand binding events.(33, 34)

Despite the above advances, a longstanding general drawback of photo-CIDNP is the limited subset of nuclei that show appreciable hyperfine couplings to the unpaired electrons within the radical pair.(12, 21, 35) Thus, nuclei lacking hyperfine couplings remain LC-photo-CIDNP-invisible at low concentration, and the extent of attainable structural and dynamic information remains limited. Representative examples targeting the ^1^H-detected ^13^C LC-photo-CIDNP 1D and 2D spectra of uniformly ^1^H,^13^C-labeled Trp and the SH3 protein (i.e., the N-terminal domain of the SH3 protein from *Drosophila melanogaster*), respectively, are shown in Fig. 1C.

A potential solution is to propagate LC-photo-CIDNP-enhanced magnetization from directly polarized nuclei to their scalar-coupled, non-photo-CIDNP-active neighbors via COSY. This well-known NMR experiment (36) targets scalar-coupling-mediated magnetization transfer. On the other hand, COSY is restricted to directly coupled nearby spins within the weak-coupling regime and relies on the generation of antiphase coherences. The latter are susceptible to cancellation effects if the extent of scalar coupling is comparable to spectral linewidths.(36–38) In contrast, Total Correlation Spectroscopy (TOCSY), also known as Homonuclear Hartmann-Hahn spectroscopy (HOHAHA), is a well-established NMR technique that does not suffer from the above shortcomings.(39–41) TOCSY employs an isotropic mixing scheme to efficiently transfer in-phase magnetization across multiple bonds within a scalar-coupled network via strong scalar-coupling Hamiltonian.(36, 38, 40) During the isotropic mixing period, a suitably designed radiofrequency (RF) field effectively eliminates chemical-shift terms, thereby maintaining the Hartmann–Hahn matching condition across the homonuclear spin network (Fig. 1D).(36, 39, 40) This feature renders TOCSY particularly attractive to extend LC-photo-CIDNP enhancements beyond the subset of directly hyperpolarized nuclei.

Prior studies exploring through-bond ^1^H photo-CIDNP magnetization transfer focused on high-concentration samples (mM regime). These investigations examined 1D COSY-based approaches on free tryptophan (Trp) and proteins.(42, 43) The COSY transfer, however, is not ideal to propagate magnetization beyond vicinal couplings. TOCSY-mediated ^1^H photo-CIDNP polarization transfer was also carried out.(43) The latter work, however, was exclusively based on homonuclear 1D ¹H experiments and targeted only solvent-exposed resonances at high (2 mM) sample concentration. Moreover, the isotropic mixing sequence (MLEV-17) employed in this study has limited bandwidth capabilities and requires higher RF power than more recently developed isotropic-mixing schemes including DIPSI-2/3, FLOPSY-16.(39, 44, 45)

The lack of multidimensional and heteronuclear approaches in these earlier studies limits their impact, especially in light of the prominent role of heteronuclear NMR in modern spectroscopy, where ^1^H- and ^13^C-detected experiments provide essential structural and dynamic information.(36, 46, 47) In all, integration of LC-photo-CIDNP with modern isotropic mixing sequences and exploration of both ^1^H and ^13^C TOCSY remain fully unexplored. Moreover, TOCSY through-bond polarization transfer experiments on proteins at low μM concentration have never been performed to date.

Here, we address the above gap of knowledge by synergistically combining LC-photo-CIDNP with two highly efficient isotropic TOCSY mixing schemes, DIPSI-2 and FLOPSY-16.(39, 44, 45) First, we demonstrate that ^1^H-and ^13^C-detected TOCSY LC-photo-CIDNP of Trp efficiently transfers hyperpolarization from the directly polarized nuclei to several scalar-coupled nearby spins. The TOCSY-enhanced resonances pertain to ^1^H^α^ and, remarkably, also to carbonyl carbons (^13^C’). Importantly, both nuclei are diagnostic of protein secondary structure.(48, 49) Notably, before the advances presented here, ^1^H^α^ polarization could only be detected via ^13^C LC-photo-CIDNP followed by reverse-INEPT and ^1^H detection (in the absence of TOCSY), thus requiring ^13^C isotopic enrichment. In contrast, the ^1^H TOCSY LC-photo-CIDNP approach developed in this work can also be applied to biomolecules at natural-abundance, and enables detection of resonance that remain invisible via conventional LC-photo-CIDNP.(47, 50) In the case of experiments targeting ^13^C TOCSY transfer to carbonyl carbons (^13^C’), we show hyperpolarization of these resonances for the first time. Second, we extend our hyperpolarization transfer strategies to a model protein, and show that 1D ^13^C-detected, 1D and 2D ^1^H-detected-^13^C-TOCSY LC-photo-CIDNP reveals previously unobservable aromatic and aliphatic side-chain resonances within minutes, down to 10 μM concentration.

In summary, this work expands the achievable LC-photo-CIDNP hyperpolarization to additional through-bond coupled ^1^H and ^13^C amino-acid and protein resonances. In this way, uniform detectability is approached at low-μM sample concentration.

## Results and Discussion

### Experimental design

Unfortunately, LC-photo-CIDNP does not typically provide uniform sensitivity enhancements across the spin network of target molecules. This outcome is perhaps not surprising, given that both geminate and F-pair polarizations are expected to scale with the hyperfine coupling constant of individual nuclei in solution.(22, 23) This undesirable feature severely attenuates the extent of attainable sensitivity enhancements, given that only a few selected nuclei benefit from photo-CIDNP polarization while others remain unenhanced, hence typically undetected (Fig. 1C).

To overcome the above limitation, we reasoned that transiently gained hyperpolarization of selected resonances could be combined with through-bond TOCSY-type isotropic mixing.(39–41) As a result, the LC-photo-CIDNP polarization generated within a dilute NMR sample could get propagated throughout the scalar-coupled spin network of the target molecule. This concept is pictorially illustrated in Fig. 1D. Only when isotropic mixing is applied immediately after hyperpolarization, LC-photo-CIDNP enhanced *I_1_* polarization experiences through-bond hyperpolarization transfer to *I_2_* in the presence of *J_12_* scalar coupling. This process increases the observable *I_2_* intensity, thereby expanding the sensitivity enhancement beyond the directly hyperpolarized resonances.

Key equations underlying the TOCSY transfer are listed in Fig. 1D, based on a simplified model comprising two J-coupled nuclear spins. These relations illustrate how polarization initially localized on one spin (*I_1_*) can be transferred to another (*I_2_*) via the *J_12_* scalar coupling during isotropic mixing. The time evolution of longitudinal magnetization during isotropic mixing can be described by terms that depend on the enhancement factor *ε*, the scalar coupling constant *J_12_* and the TOCSY mixing time. Properly optimized isotropic mixing times (*τ_mix_*) enable hyperpolarization to be redistributed while minimizing relaxation losses, thereby improving detectability of otherwise weak or undetectable resonances. The section below highlight how adoption of TOCSY strategy leads to significant increases in spectral coverage, enabling more comprehensive and widespread hyperpolarization enhancements in dilute solution.

### 1H TOCSY propagates 1H LC-photo-CIDNP hyperpolarization to nearby resonances

TOCSY-mediated ^1^H hyperpolarization transfer was first evaluated on natural-abundance Trp, a readily available model compound. In LC-photo-CIDNP, only selected resonances are directly enhanced (Fig. 2A). On the other hand, upon inclusion of TOCSY isotropic mixing (rf field = 10,000 Hz / τ_mix_ = 50 ms), additional ^1^H resonances experience significant intensity enhancements (Fig. 2A, see green arrows). This result demonstrates that TOCSY can propagate LC-photo-CIDNP hyperpolarization through scalar-coupled ^1^H nuclear-spin networks. To quantitatively assess the TOCSY-mediated LC-photo-CIDNP hyperpolarization, enhancement factor (ε) were assessed according to

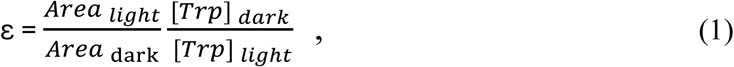

**Figure 2.**
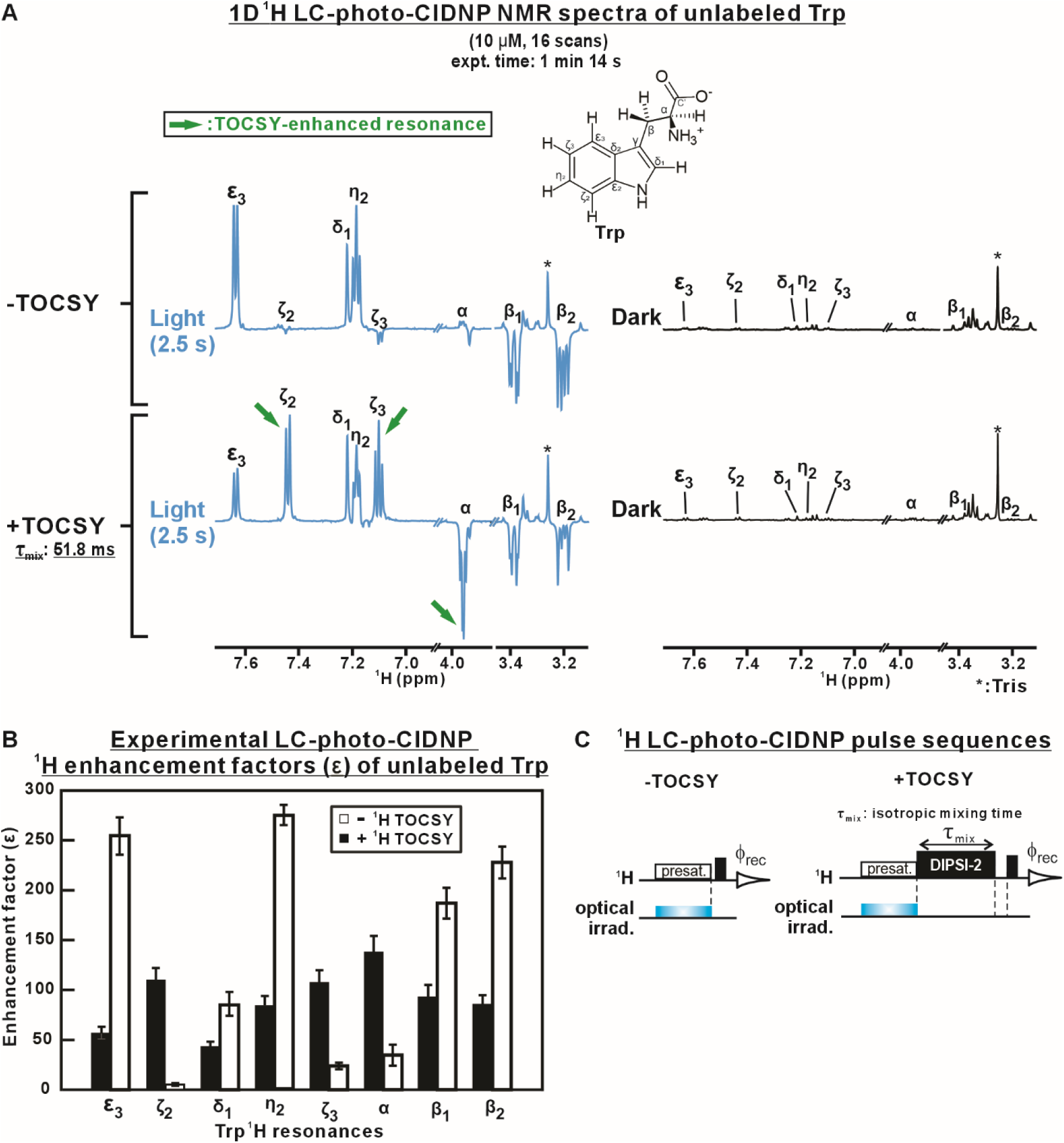
^1^H TOCSY-mediated hyperpolarization transfer enables uniform enhancement of unlabeled-Trp resonances. *(A)* Representative ^1^H LC-photo-CIDNP spectra of unlabeled Trp acquired in the absence and presence of TOCSY isotropic mixing (rf field strength = 10,000 Hz / τ_mix_ = 50 ms). In the absence of TOCSY, photo-CIDNP selectively enhances aromatic resonances. In the presence of TOCSY, the initial hyperpolarization gets redistributed across the nearby scalar-coupled nuclei, resulting in the substantial enhancement of previously poorly hyperpolarized or undetectable resonances (green arrows), including ζ₂, ζ₃, and α. Dark spectra are shown as controls. *(B)* Experimental enhancement factors (ε) of individual ^1^H resonances of unlabeled Trp in the absence and presence of TOCSY. While directly hyperpolarized aromatic resonances exhibit reduced enhancement due to polarization redistribution, substantial enhancements are observed for initially poorly polarized resonances, demonstrating efficient hyperpolarization transfer. All data are shown as avg. ± SE (n = 2). *(C)* Pulse sequences used for ^1^H LC-photo-CIDNP (left) and TOCSY-mediated LC-photo-CIDNP (right). The presat. abbreviation denotes solvent presaturation. All samples were in 10 mM potassium phosphate (pH 7.2). Data were collected in the presence of the fluorescein photosensitizer (10 μM), oxygen-scavenging enzymes and vitamin C. See Materials and Methods for details.

where *Area* denotes the integrated Trp ¹H-resonance area and [Trp] is the Trp concentration, under LED-on (light) and LED-off (dark) conditions. Numerical values of the enhancement factors (ε) of individual ^1^H resonances are shown in Table 1. As expected, the directly hyperpolarized resonances show reduced enhancements in the presence of TOCSY, consistent with redistribution of the initially gained polarization. In contrast, non-of weakly hyperpolarized resonances displayed substantial enhancement in the presence of TOCSY, due to the presence of efficient through-bond hyperpolarization transfer. The above results demonstrate that TOCSY is capable of efficiently redistributing localized LC-photo-CIDNP polarization throughout scalar-coupled spin networks, thus enabling detection of resonances that are not directly LC-photo-CIDNP-active. The pulse sequences employed in these experiments are shown in Fig. 2*C*. Given that photo-CIDNP directly generates non-equilibrium longitudinal magnetization, no coherence-generating excitation pulse is required prior to isotropic mixing. The DIPSI-2 mixing sequence efficiently transfers longitudinal polarization throughout the scalar-coupled spin network.

**Table 1.** Enhancement factors for LC-photo-CIDNP hyperpolarization of unlabeled Trp ^1^H resonances in the absence and presence of TOCSY.

| $^1\text{H}$<br>resonance | Enhancement factor ($\epsilon$) | |
| --- | --- | --- |
|  | -TOCSY | +TOCSY |
| $\epsilon 3$ | $255 \pm 18.9$ | $57 \pm 6.2$ |
| $\zeta 2$ | $6 \pm 0.6$ | $110 \pm 11.6$ |
| $\delta 1$ | $86 \pm 11.8$ | $44 \pm 4.6$ |
| $\eta 2$ | $275 \pm 10.9$ | $85 \pm 9.4$ |
| $\zeta 3$ | $24 \pm 3.1$ | $108 \pm 11.9$ |
| $\alpha$ | $35 \pm 10.5$ | $139 \pm 15.6$ |
| $\beta 1$ | $187 \pm 15.6$ | $93 \pm 11.2$ |
| $\beta 2$ | $228 \pm 15.9$ | $86 \pm 9.0$ |

The phases of the TOCSY-enhanced LC-photo-CIDNP resonances are diagnostic of the polarization-transfer pathways within the Trp spin system. The ^1^H^α^ resonance exhibits the same negative phase as the ^1^H^β^ resonances, consistent with polarization transfer occurring from the ^1^H^β^ spins via scalar couplings. Similarly, the phases of the ^1^H^ζ2^ and ^1^H^ζ3^ resonances are consistent with polarization transfer originating from neighboring aromatic resonances. At the relatively short mixing time employed here (51.8 ms), polarization transfer occurs predominantly within the respective aliphatic and aromatic spin networks, with only limited transfers among these regions. This effective separation minimizes interference between polarization-transfer pathways originating from the two distinct spin networks.

### 13C TOCSY coupled with ^13^C LC-photo-CIDNP extends hyperpolarization to carbonyl-carbon and other resonances

Next, we extended our strategy discussed to the transfer of ^13^C hyperpolarization. Although conventional ^1^H LC-photo-CIDNP provides high sensitivity, the limited chemical-shift dispersion of ^1^H nuclei often result in severe spectral overlaps. In contrast, ^13^C’s much wider chemical-shift dispersion leads to substantially improved spectral resolution. This feature renders the ^13^C nucleus particularly advantageous to the analysis of complex biomolecules. The LC-photo-CIDNP enhancement factors ε of individual ^13^C resonances in the absence and presence of TOCSY are shown in Fig. 3B and Table 2. When TOCSY isotropic mixing is absent, LC-photo-CIDNP selectively enhances only a limited subset of ^13^C resonances of uniformly ^13^C,^15^N-labeled Trp (Trp-U-^13^C, ^15^N, Fig. 3A, upper panel). In contrast, following incorporation of a 10 ms FLOPSY-16 mixing (rf field strength = 16,666 Hz), several additional ^13^C resonances become detectable, including carbonyl (C′), aromatic (*ζ*_2_, *ζ*_3_), and aliphatic (*β*) carbons (Fig. 3A, lower panel). The newly observed resonances are highlighted by green arrows. These data show the presence of efficient redistribution of the initially generated ^13^C LC-photo-CIDNP polarization throughout the ^13^C spin network during isotropic mixing. As expected, the degree of polarization redistribution depends on the TOCSY mixing time (SI Appendix Fig. S1A).

**Figure 3.**
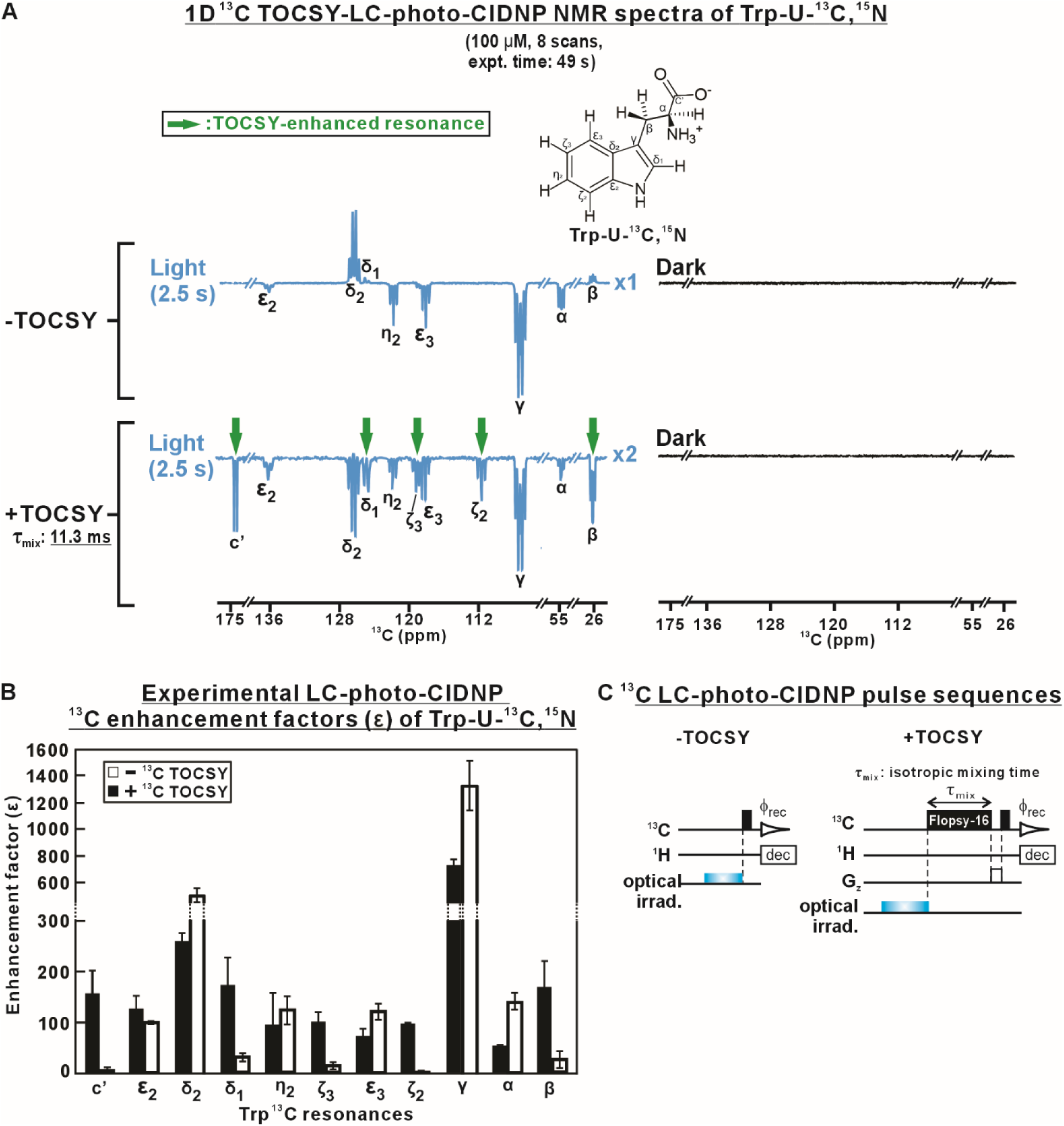
^13^C TOCSY enables uniform propagation of hyperpolarization across all resonances of ^13^C-labeled Trp. *(A)* Representative ^13^C LC-photo-CIDNP spectra of Trp-U-^13^C-^15^N acquired in the presence of TOCSY (rf field strength = 16,666Hz, τ_mix_ = 10 ms). TOCSY redistributes the initial hyperpolarization throughout the network of J-coupled nuclei, resulting in the appearance and substantial enhancement of previously poorly hyperpolarized or undetectable resonances (green arrows), including ^13^C^β^, ^13^C^ζ₂^, ^13^C^η2^, ^13^C^δ1^ and ^13^C’. Dark (i.e., LED-off) spectra are shown as controls. *(B)* Experimental enhancement factors (ε) of individual ^13^C resonances of Trp-U-^13^C, ^15^N in the absence and presence of TOCSY. All data are shown as avg. ± SE (n = 2). *(C)* Pulse sequences used for ^13^C LC-photo-CIDNP (left) and TOCSY-enhanced LC-photo-CIDNP (right) experiments. A FLOPSY-16 isotropic mixing period was inserted before acquisition to transfer LC-photo-CIDNP hyperpolarization throughout the J-coupled spin network. Data were collected in 10 mM potassium phosphate (pH ∼7.2) in the presence of the fluorescein photosensitizer (25 μM), oxygen-scavenging enzymes and vitamin C. See Materials and Methods for details.

**Table 2.**
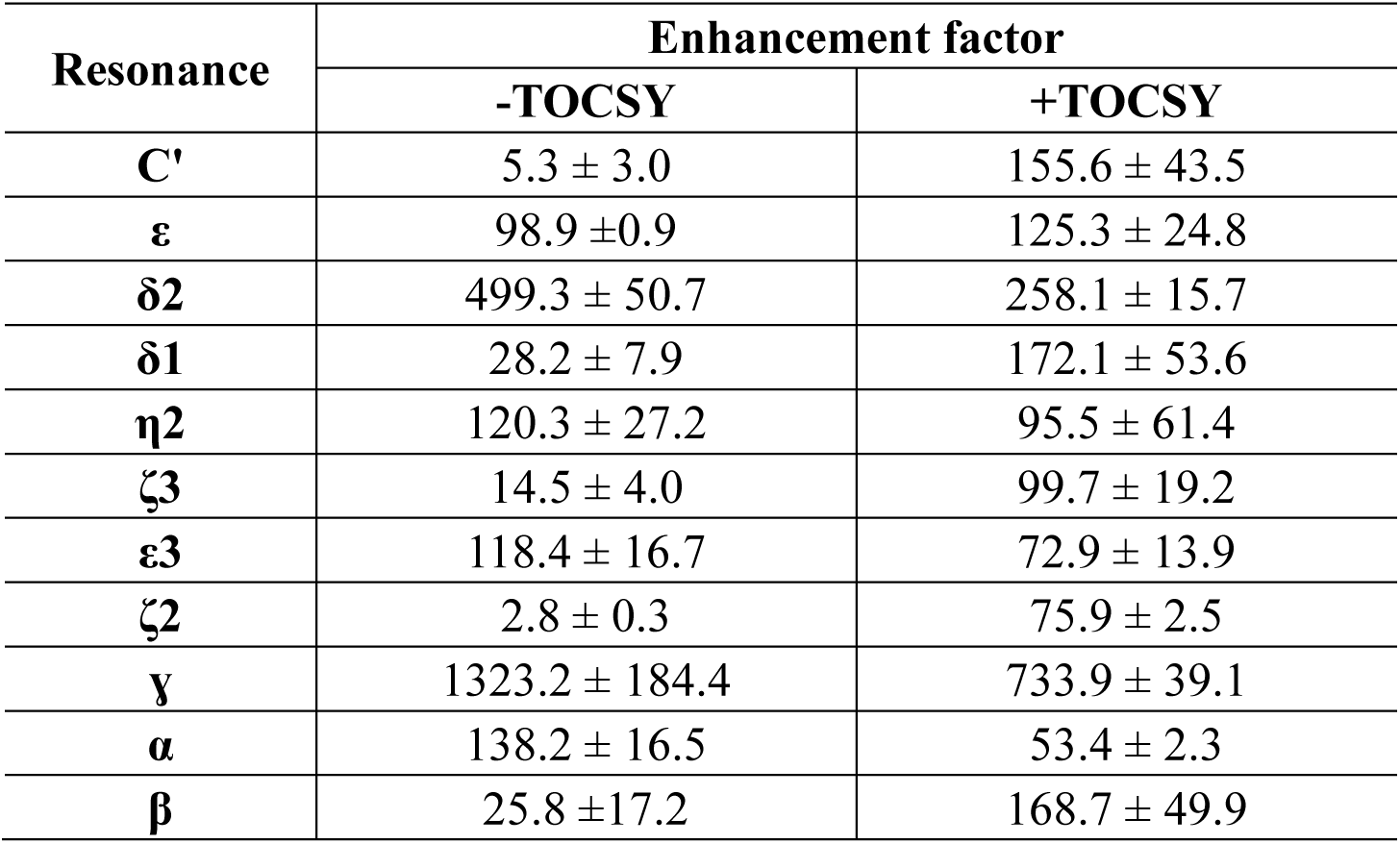
Enhancement factors for LC-photo-CIDNP hyperpolarization of ^13^C resonances of Trp-U-^13^C, ^15^N in the absence and presence of TOCSY.

| Resonance | Enhancement factor |  |
| --- | --- | --- |
|  | -TOCSY | +TOCSY |
| **C'** | $5.3 \pm 3.0$ | $155.6 \pm 43.5$ |
| **$\epsilon$** | $98.9 \pm 0.9$ | $125.3 \pm 24.8$ |
| **$\delta 2$** | $499.3 \pm 50.7$ | $258.1 \pm 15.7$ |
| **$\delta 1$** | $28.2 \pm 7.9$ | $172.1 \pm 53.6$ |
| **$\eta 2$** | $120.3 \pm 27.2$ | $95.5 \pm 61.4$ |
| **$\zeta 3$** | $14.5 \pm 4.0$ | $99.7 \pm 19.2$ |
| **$\epsilon 3$** | $118.4 \pm 16.7$ | $72.9 \pm 13.9$ |
| **$\zeta 2$** | $2.8 \pm 0.3$ | $75.9 \pm 2.5$ |
| **$\gamma$** | $1323.2 \pm 184.4$ | $733.9 \pm 39.1$ |
| **$\alpha$** | $138.2 \pm 16.5$ | $53.4 \pm 2.3$ |
| **$\beta$** | $25.8 \pm 17.2$ | $168.7 \pm 49.9$ |

The ability to detect backbone carbonyl (C′) resonances at high sensitivity is expected to be particularly valuable in the case of intrinsically disordered proteins (IDPs) or protein regions (IDRs). The structural characterization of these biomolecules is often hindered by spectral overlaps due to the absence of persistent secondary and(or) tertiary structure. Backbone carbonyl resonances are distributed over a much wider chemical-shift range and are highly sensitive to local backbone conformation. Therefore, transferring of LC-photo-CIDNP polarization to carbonyl (^13^C’) and other ^13^C nuclei enables extending the structural information accessible by LC-photo-CIDNP alone, facilitating residue-specific characterization of conformational heterogeneity and transient structural elements. The pulse sequences used pertaining to these experiments are shown in Fig. 3*C*.

As in the case of DIPSI-2, the augmented non-equilibrium longitudinal magnetization generated via LC-photo-CIDNP can be directly fed to FLOPSY-16 mixing, with no need for a coherence-generating excitation pulse. Given the broadband isotropic mixing characteristics of FLOPSY-16, this pulse train efficiently redistributes polarization throughout the scalar-coupled spin network of the LC-photo-CIDNP-enhanced resonances, thus extending hyperpolarization from directly polarized sites to indirectly polarized nuclei across the broad ^13^C chemical-shift range. It is worth noting that, among the ^13^C LC-photo-CIDNP resonances, only ^13^C^δ2^ exhibits a positive phase, while the remaining resonances are negative. This behavior likely arises from the combined contribution of polarization transfer via multiple pathways within the scalar-coupled spin network. It is expected that transfers of polarizations with opposite signs to a given single nucleus lead to partial cancellation effects. As a result, redistribution of the initially localized hyperpolarization throughout an extended spin network may result in an overall reduction in signal-to-noise ratio, due to the competing transfer pathways. Regardless, TOCSY-mediated polarization transfer enables detection of multiple resonances that are not directly hyperpolarized via LC-photo-CIDNP. This effect substantially increases the number of observable resonances and expands the residue-specific information accessible from the originally hyperpolarized nuclei.

### 1H-detected ^13^C photo-CIDNP coupled with TOCSY also generates extensive hyperpolarization propagation

To further improve the applicability of LC-photo-CIDNP to a wider number of resonances, we combined ^13^C hyperpolarization, reverse INEPT-based ^1^H detection and ^13^C TOCSY. Given that the gyromagnetic ratio and intrinsic sensitivity of ^1^H are substantially higher than those of ^13^C, transferring hyperpolarized ^13^C magnetization to directly bonded protons provides additional gains in sensitivity, while retaining the site selectivity of ^13^C photo-CIDNP. This strategy is particularly effective to improve the sensitivity of LC-photo-CIDNP hyperpolarization at amino-acid sites (e.g., the ^1^H^α^-^13^C^α^ pair) that experience weak ^1^H but high ^13^C hyperfine couplings within the radical pair.(7, 12, 28) As illustrated in the upper panels of Fig. 4A, conventional ^13^C RASPRINT, an ^1^H-detected ^13^C photo-CIDNP pulse sequence,(7) generates ^13^C hyperpolarization followed by transfer to the covalently linked ^1^H nuclei through via reverse INEPT. Enhancement factors of the ε_3_, ζ_2_, δ_1_, η_2_ and ζ_3_ nuclei were 223.8 ± 33.3, 7.7 ± 0.3, 63.1 ± 1.3, 153.4 ± 17.3 and 26.7 ± 15.0, respectively. To further extend the observable detectability to a wider spin network, the FLOPSY-16 isotropic mixing block(39) was incorporated immediately after photo-CIDNP hyperpolarization (Fig. 4A). As a result, the ^1^H resonances covalently linked to the TOCSY-mediated indirectly polarized ^13^Cs become detectable at substantially increased sensitivity. Specifically, the ^1^H^δ1^ resonance exhibits a dramatic enhancement factor of 664.1 ± 14.6, while the remaining ^1^H resonances also show appreciable sensitivity gains (Fig. 4B). Further, the extent of polarization redistribution can be controlled by *ad hoc-*variations in the TOCSY mixing time (during FLOPSY-16), resulting in modulations in relative spectral intensities (SI Appendix Fig. S1B). As shown in Fig. S1B, variations in FLOPSY-16 mixing time are expressed as variable numbers of isotropic-mixing loops. Introduction of a short 7.5 ms mixing period (one loop) produces a highly non-uniform redistribution of polarization. Conversely, increasing the isotropic-mixing time progressively redistributes polarization throughout the coupled ¹³C spin network, resulting in a more uniform enhancement pattern and a larger number of observable resonances.

**Figure 4.**
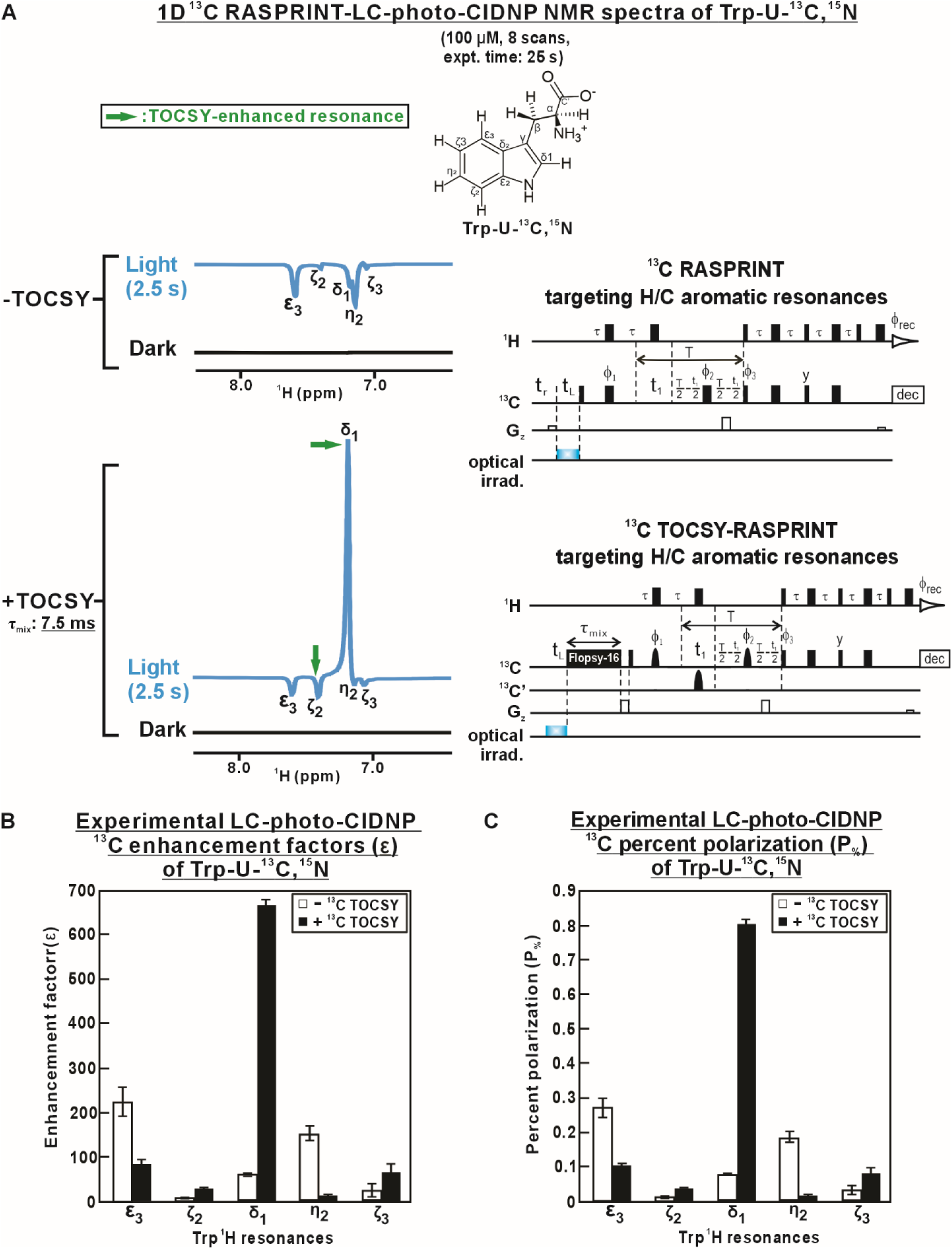
^1^H detection of ^13^C TOCSY-mediated ^13^C hyperpolarization transfer enables uniform detection of the entire ^13^C Trp spin system. *(A)* Representative ^1^H-detected ^13^C RASPRINT LC-photo-CIDNP spectra of Trp-U-^13^C-^15^N acquired in the absence and presence of ^13^C TOCSY isotropic mixing (τ_mix_ = 7.5 ms). An effective rf field strength was 6,250 Hz was employed. FLOPSY-16 isotropic mixing redistributes the initial ^13^C hyperpolarization throughout the J-coupled ^13^C spin network, augmenting detectability of previously weak or undetectable ^13^C resonances (green arrows). The pulse sequences for ^13^C RASPRINT and ^13^C TOCSY-RASPRINT are shown on the righthand side. *(B)* Experimental ^13^C LC-photo-CIDNP enhancement factors (ε) of individual ^1^H-detected ^13^C resonances of Trp-U-^13^C, ^15^N in the absence and presence of ^13^C TOCSY. *(C)* Experimental LC-photo-CIDNP percent polarization (P_%_) of individual ^13^C resonances. All data are shown as avg. ± SE (n = 2). All samples included 10 mM potassium phosphate (pH ∼7.2), 25 μM fluorescein, oxygen-scavenging enzymes and vitamin C. See Materials and Methods for details.

To better estimate absolute nuclear polarization, the enhancement factor (ε) was converted to percent polarization *P*_%_, as shown if Fig 4*C*. The percent polarization (*P*_%_) of the ^13^C aromatic resonances was derived from the enhancement factor ε according to

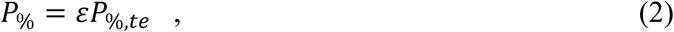

where *P*_%,*te*_ denoted the thermal-equilibrium polarization, defined as

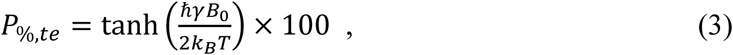

where *γ*is the gyromagnetic ratio (*γ*_13_*_C_* = 10.705 MHz/T and *γ*_1_*_H_* = 42.576 MHz/T), *B*_0_ is the applied magnetic field, and T is the absolute temperature (in K). Notably, the H^δ1^ resonance has a *P*_%_ of 0.8%, corresponding to an exceptionally high value relative to the other ^13^C sites (Fig. 4C).

### TOCSY-mediated ^13^C hyperpolarization transfer is highly effective on proteins

Next, ^13^C TOCSY-mediated LC-photo-CIDNP hyperpolarization transfer was applied to a model protein, ^13^C-labeled SH3. As shown in the upper SH3 spectrum of Fig. 5, conventional 1D LC-photo-CIDNP leads to appreciable ^13^C photo-CIDNP hyperpolarization. The enhancement of polarization, however, is confined to a limited number of resonances. Incorporation of TOCSY isotropic mixing into the pulse sequence leads to propagation of the initial hyperpolarization, resulting into clear detection of additional resonances that were absent or weak in conventional ^13^C LC-photo-CIDNP. Remarkably, increasing the isotropic mixing time leads to further redistribution of photo-CIDNP derived hyperpolarization, and enables detection of ^13^C resonances across the SH3 Trp and Tyr spin systems (Fig. 5, SI Appendix Fig. S2). Comparison between U-^13^C, ^15^N Trp and the ^13^C-labeled SH3 protein shows similar polarization transfer profiles over the entire ^13^C spectra.

**Figure 5.**
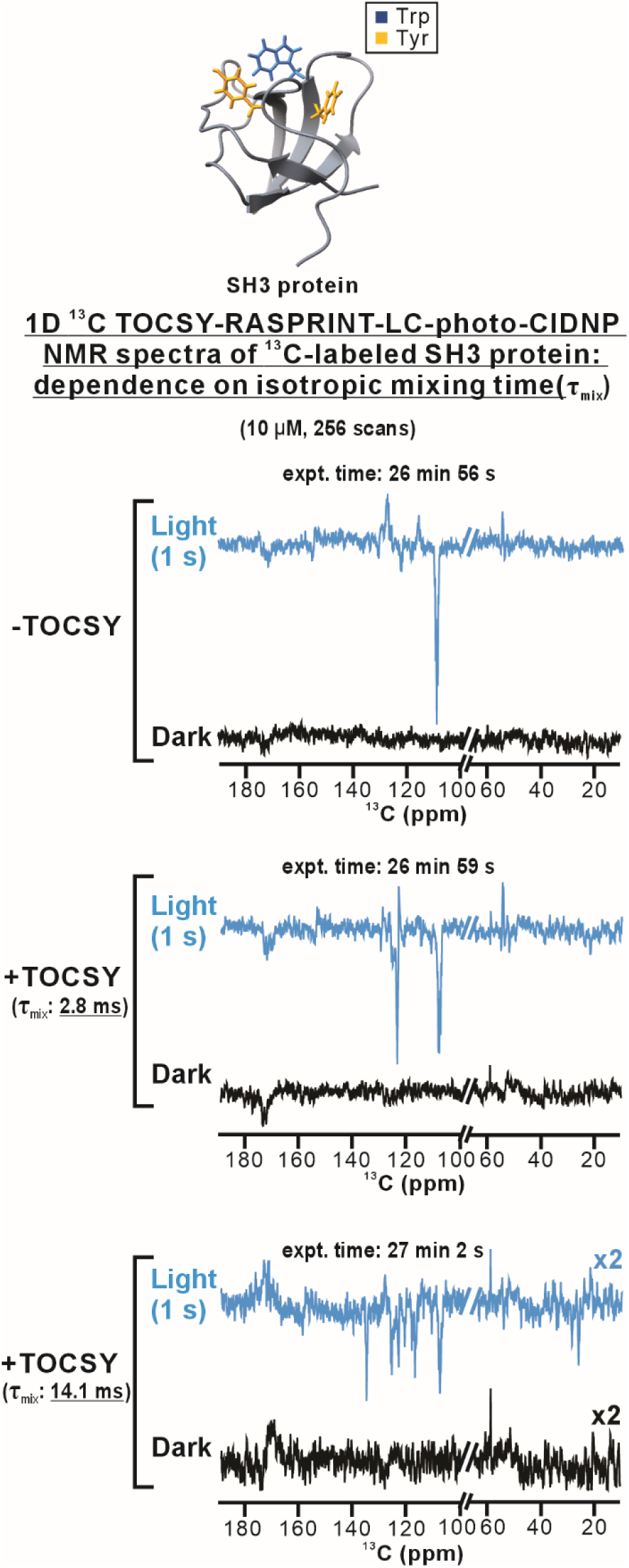
TOCSY-mediated hyperpolarization transfer enables detection of additional ^13^C resonances of the SH3 protein. Representative ^13^C TOCSY LC-photo-CIDNP spectra of ^13^C-labeled SH3 protein acquired with different TOCSY isotropic mixing times. An effective rf field strength of 16.666 Hz with short isotropic mixing (τ_mix_ = 2.8 ms) produces efficient polarization transfer to nearby J-coupled carbons while largely preserving the initial polarization. Increasing the mixing time (τ_mix_ = 14.1 ms) further redistributes the initial hyperpolarization, enabling observation of enhanced ^13^C resonances. Relevant resonance assignments are provided in Fig. S2. The 10 μM SH3 samples were in 10 mM potassium phosphate (pH ∼7.2) and included 10 μM fluorescein photosensitizer, oxygen-scavenging enzymes and vitamin C. See Materials and Methods for details.

In all, the above data show that TOCSY-mediated redistribution of LC-photo-CIDNP hyperpolarization is highly effective across both free amino acids and proteins. Further, the protein enhancements conveniently apply to both the Trp and Tyr residues. This effect is likely contributed by well-known intramolecular transient Trp-to-Tyr electron-transfer processes within protein frameworks.(51, 52) In turn, this phenomenon leads to a competitive advantage for the LC-photo-CIDNP hyperpolarization of proteins carrying Trp and Tyr, over hyperpolarization of the individual amino acids, which typically requires distinct photosensitizer dyes, at low concentration. Together, the above capabilities extend the utility of photo-CIDNP beyond signal enhancement and establish ^13^C TOCSY-photo-CIDNP as a versatile platform for structural and dynamic investigations of complex biomolecules.

### 1D and 2D ^1^H-detected ^13^C TOCSY-RASPRINT further expands hyperpolarization across the SH3-protein spin network

In order to maximize experimental sensitivity and resolution for the NMR detection of proteins, ^13^C photo-CIDNP hyperpolarization was combined with ^1^H detection. The SH3 model protein was employed in both 1D and 2D experiments. Conventional ^1^H-detected ^13^C RASPRINT in the presence of fluorescein leads to only a subset of directly hyperpolarized Trp and Tyr resonances(16), as shown in Fig. 6A-B. The LC-photo-CIDNP phases of Trp and Tyr are opposite, likely due to variations in g factors within the transient radical pairs of these amino acids.(16) All Trp aromatic resonances are emissive (negative phase), while the two Tyr aromatic resonances display opposite phase, one being absorptive (positive) and the other emissive (negative). Trp and Tyr residues can be readily distinguished based on their opposite signal phases. The 2D experiment provides substantially improved spectral resolution, and the corresponding resonance assignments(53) are shown in Fig. 6C. Although LC-photo-CIDNP provides a substantial gain in sensitivity, it does not enable detection of all resonances, even in 2D experiments, due to site selective enhancement or resonances belonging to redox-active amino acids. On the other hand, ^13^C TOCSY-RASPRINT alleviates this feature by enhancing additional Trp and Tyr resonances in 1D and 2D spectra (Fig. 6B-D). As expected in the case of proteins, a relatively short mixing time is ideal to mediate polarization transfer through the one-bond ^13^C–^13^C couplings of SH3. Efficient polarization transfer from the intense W36_N_ and W36_U_ ^13^Cγ resonances to the directly bonded Cδ1 nuclei results in a prominent resonance at approximately 7.2 ppm in both the 1D and 2D, ^13^C TOCSY-RASPRINT spectra of SH3 (Fig 6*B* and *D* upper panels). While longer mixing times further improved resonance coverage by transferring polarization to more distant ^13^C nuclei, although some directly hyperpolarized resonances decrease in intensity due to polarization redistribution (Fig 6B and D lower panel).

**Figure 6.**
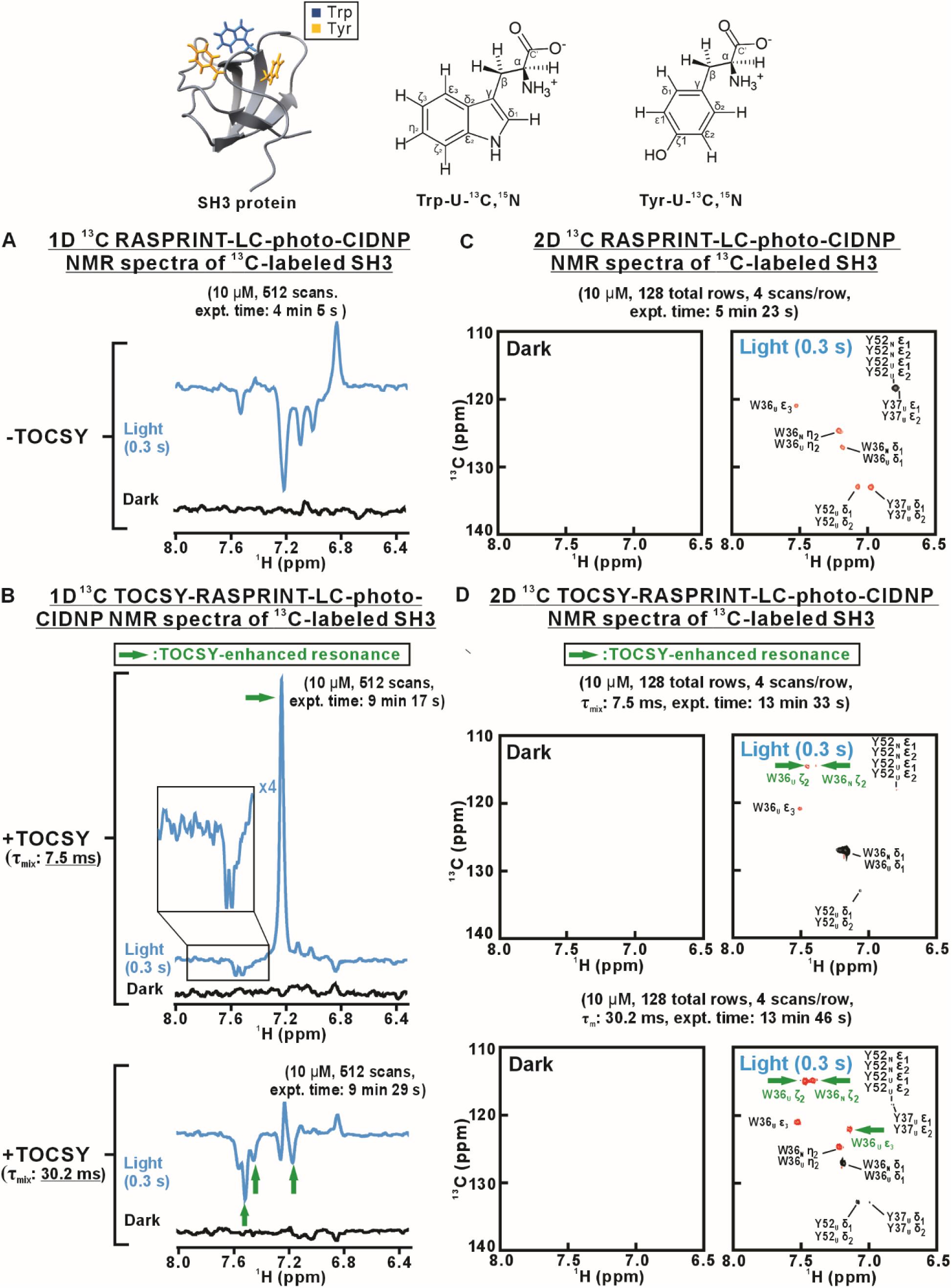
^1^H-detected ^13^C TOCSY-RASPRINT enables augmented detection of hyperpolarized resonances of the SH3 protein. *(A)* Representative 1D ^1^H-detected ^13^C RASPRINT LC-photo-CIDNP spectra of ^13^C-labeled SH3 protein acquired in the absence of TOCSY. Under light (LED-on) conditions, only directly hyperpolarized Trp and Tyr resonances are detected. *(B)* Representative 1D ^13^C TOCSY-RASPRINT LC-photo-CIDNP spectra acquired with isotropic mixing times of 7.5 and 30.2 ms, with an effective rf field strength of 6,250 Hz. Enhanced resonances arising from polarization transfer are highlighted in green. *(C)* 2D ^13^C RASPRINT spectra acquired in the presence of TOCSY. Under light conditions, only directly hyperpolarized Trp and Tyr resonances are observed*. (D)* 2D ^13^C TOCSY-RASPRINT spectra of SH3 protein with isotropic mixing times of 7.5 and 30.2 ms. Enhanced resonances arising from polarization transfer are highlighted in green. The 10 μM SH3 samples were in 10 mM potassium phosphate (pH ∼7.2), and contained 10 μM fluorescein, oxygen-scavenging enzymes and vitamin C. See Materials and Methods for details.

Taken together, the 1D and 2D protein data demonstrate that TOCSY-RASPRINT is an effective tool to redistribute localized photo-CIDNP hyperpolarization, thereby substantially increasing the number of NMR-observable hyperpolarized resonances at low concentrations in solution. In all, this technology extends the information content of both 1D and 2D LC-photo-CIDNP experiments on proteins in dilute solution.

### Conclusions

In this work, we show that optical irradiation followed by radiofrequency isotropic-mixing schemes provide an effective strategy to overcome the intrinsic site-selectivity of LC-photo-CIDNP. By redistributing LC-photo-CIDNP-derived hyperpolarization through scalar-coupled spin networks, TOCSY enables the detection of resonances that are inaccessible via conventional LC-photo-CIDNP, thereby substantially expanding spectral coverage. The applicability of the above strategy to both free amino acids and the SH3 model protein highlights the fact that this technology applies to a broad range of molecular sizes. In summary, our results establish TOCSY-assisted LC-photo-CIDNP as a practical platform for sensitive heteronuclear-correlation multidimensional NMR spectroscopy to investigate biomolecular conformation in dilute solution. We anticipate that the strategy highlighted here will substantially broaden the information content of LC-photo-CIDNP by enabling widespread hyperpolarization across many resonances of dilute biomolecules.

## Materials and Methods

### Materials

Uniformly ^13^C, ^15^N-labeled tryptophan (Trp-U-^13^C, ^15^N) was obtained from Cambridge Isotope Laboratories (Tewksbury, MA, USA). Fluorescein sodium salt, vitamin C (ascorbic acid, VC), and the oxygen-scavenging system comprising glucose oxidase from *Aspergillus niger* (GO, EC 1.1.3.4, lyophilized powder) and catalase from bovine liver (CAT, EC 1.11.1.6, lyophilized powder) were purchased from Millipore Sigma. In addition, D-glucose (U-D_12_) and D-Glucose (U-¹³C₆) were purchased from Cambridge Isotopes.

### SH3 protein preparation

The uniformly ^13^C-labeled SH3 protein was expressed and purified as described.(54) Briefly, *E. coli* BL21 cells were grown in M9 minimal medium supplemented with ^13^C sources (D-Glucose-^13^C_6_). Protein expression was induced with isopropyl β-D-1-thiogalactopyranoside (IPTG) at an OD600 of approximately 0.6, and cells were harvested after further growth to an optical density at 600 nm (OD600) greater than 1.2. Following cell lysis, the protein was purified by anion-exchange chromatography and size-exclusion chromatography. The purified protein was subsequently dialyzed against Tris buffer (50 mM Tris, 5 mM MgCl₂, 50 mM KCl, pH 7.2). The purified SH3 solution was then aliquoted, flash-frozen in liquid nitrogen, and stored at −80 °C. SH3 protein concentration was determined by UV–visible electronic absorption spectroscopy, using an extinction coefficient of 8,480 M⁻¹ cm⁻¹ at 280 nm.(7)

### LC-photo-CIDNP NMR sample preparation

All LC-photo-CIDNP NMR samples, including U-^13^C, ^15^N-Trp and ^13^C-labeled SH3, were prepared in 10 mM potassium phosphate buffer (pH 7.2) containing 5% D_2_O, 0.15 μM glucose oxidase (GO), 0.10 μM catalase (CAT), 10 μM fluorescein, and 2.0 μM vitamin C and D-Glucose (U-D_12_) (2.5 mM) was added approximately 10 min prior to NMR data acquisition. Stock solutions of GO, CAT, fluorescein, and SH3 were freshly thawed on ice immediately before use, whereas vitamin C solutions were prepared fresh on the day of each experiment. All NMR experiments were performed on an NMR spectrometer operating at 600 MHz. DSS-*d*_6_ in D_2_O was used as an external chemical-shift reference in all experiments.

### LC-photo-CIDNP data collection under dark conditions for enhancement factor determination

Enhancement factors ε of ^1^H-acquire experiments were determined with the help of a reference sample comprising 10 μM unlabeled Trp in 10 mM potassium phosphate buffer (pH 7.2). Enhancement factors of ^13^C LC-photo-CIDNP experiments were assessed with the aid of reference data collected under dark conditions on a highly concentrated Trp-U-^13^C, ^15^N sample (5 mM) in 10 mM potassium phosphate buffer (pH 7.2). Both experiments were carried out with a long recycle delay (5 s), to allow for complete longitudinal relaxation between transients.

### LED sources and optical irradiation setup

A UHP-FB-LED 450 light source (Prizmatix, Holon, Israel; peak emission at 453 nm) was coupled to a 4.1 m polymer optical fiber (POF; 1.5 mm diameter, numerical aperture = 0.5; Prizmatix) for the all LC-photo-CIDNP experiment. The optical fiber was inserted into a 4 mm NMR tube (427-PP-7, Wilmad-LabGlass, Buena, NJ, USA), which was then positioned concentrically inside a 5 mm NMR tube containing the sample. This configuration ensured reproducible alignment of the optical fiber with the NMR sample volume for efficient irradiation. The LED output power measured at the fiber tip was 450 mW. LED irradiation times for individual experiments are provided in the relevant figures.

### Experimental Conditions for LC-Photo-CIDNP experiments in the absence and presence of TOCSY

All NMR data were collected at 600 MHz on a Bruker Avance III NMR spectrometer equipped with a 5 mm ^1^H/^13^C/^15^N-^19^F triple-resonance cryogenic probe (TCI-F) equipped with z-axis gradients. LC-photo-CIDNP experiments were performed with an UHP-FB-LED 450 optical irradiation source, with light delivered to the sample through a polymer optical fiber as described above. In 1D, ^1^H LC-photo-CIDNP experiments performed in the absence of presence of TOCSY, the ^1^H carrier frequency was centered on the HDO resonance (ca. 4.7 ppm), and solvent presaturation was applied. Data were collected with 16 scans, 32,768 total points, acquisition time of 1.95 s, a recycle delay of 0.05 s, a LED irradiation time of 2.5 s and a mixing time of 51.8 ms. 1D ^13^C LC-photo-CIDNP experiments in the absence or presence of TOCSY were performed with the ^13^C carrier frequency centered at 120 ppm using 16,384 complex points, an acquisition time of 0.22 s, a recycle delay of 2.5 s, a LED irradiation time of 2.5 s, and isotropic mixing times of 11.3 ms and 8 scans. In the case of 1D ^13^C TOCSY-RASPRINT experiments targeting the aromatic region, the ^1^H and ^13^C carrier frequencies were centered at 4.7 and 120 ppm, respectively. Spectra were acquired with 8 scans, 4,096 total points, an acquisition time of 0.24 s, a recycle delay of 1 s, a LED irradiation time of 2.5 s and an isotropic mixing time of 7.5 ms.

1D, ^13^C TOCSY-CIDNP experiments on the SH3 protein had ^1^H and ^13^C carrier frequencies centered at 5.5 and 120 ppm, respectively. Data were collected with 256 scans, 16,384 total points, an acquisition time of 0.22 s, a recycle delay of 5 s, a LED irradiation time of 1 s, and isotropic mixing times of 2.8 and 14.1 ms. The 1D ^13^C TOCSY-RASPRINT experiments on the SH3 protein were carried out with the ^1^H and ^13^C carrier frequencies centered at 4.7 and 120 ppm, respectively. Spectra were acquired with 512 scans, 4,096 complex points, an acquisition time of 0.24 s, a recycle delay of 0.5 s, an LED irradiation time of 0.3 s and isotropic mixing times of 7.5 ms and 30.2 ms. The 2D ^13^C TOCSY-RASPRINT (aromatic) experiments performed on the SH3 protein included data acquisition with the ^1^H and ^13^C carrier frequencies centered at 4.7 and 120 ppm, respectively. The data matrix consisted of 4,096 complex points in the direct dimension and 128 increments in the indirect dimension, with 4 scans per increment, an acquisition time of 0.24 s, a recycle delay of 1 s, LED irradiation time of 0.3 s and isotropic mixing times of 7.5 and 30.2 ms.

1D NMR data were processed with MestReNova (version 15.1.0), while all 2D NMR data were processed with NMRPipe (v. 9.0.0-b10) and visualized with NMRDraw and NMRViewJ (v. 2009.015.15.35).(55, 56) The 1D data were processed with 4-fold zero-filling and exponential apodization (with 1 - 5 Hz line-broadening). The 2D data were processed with 2-fold zero-filling followed by unshifted cosine-bell-square apodization in both dimensions.

## Acknowledgments

We thank Heike Hofstetter for technical assistance, and members of the Cavagnero group for helpful discussions. This work was supported by the National Institutes of Health (grants R01GM125995 and R35GM161252 to SC). The Bruker Avance III 600 NMR spectrometer in the Department of Chemistry was supported by NIH grant S10 OD012245.

## Data Availability

The data presented in this article will be shared upon reasonable request to the corresponding author.

