## Supplementary Information for "Efficient Through-Bond Propagation of Nuclear-Spin Hyperpolarization via TOCSY-Enhanced LC-Photo-CIDNP"

### Supporting Figures

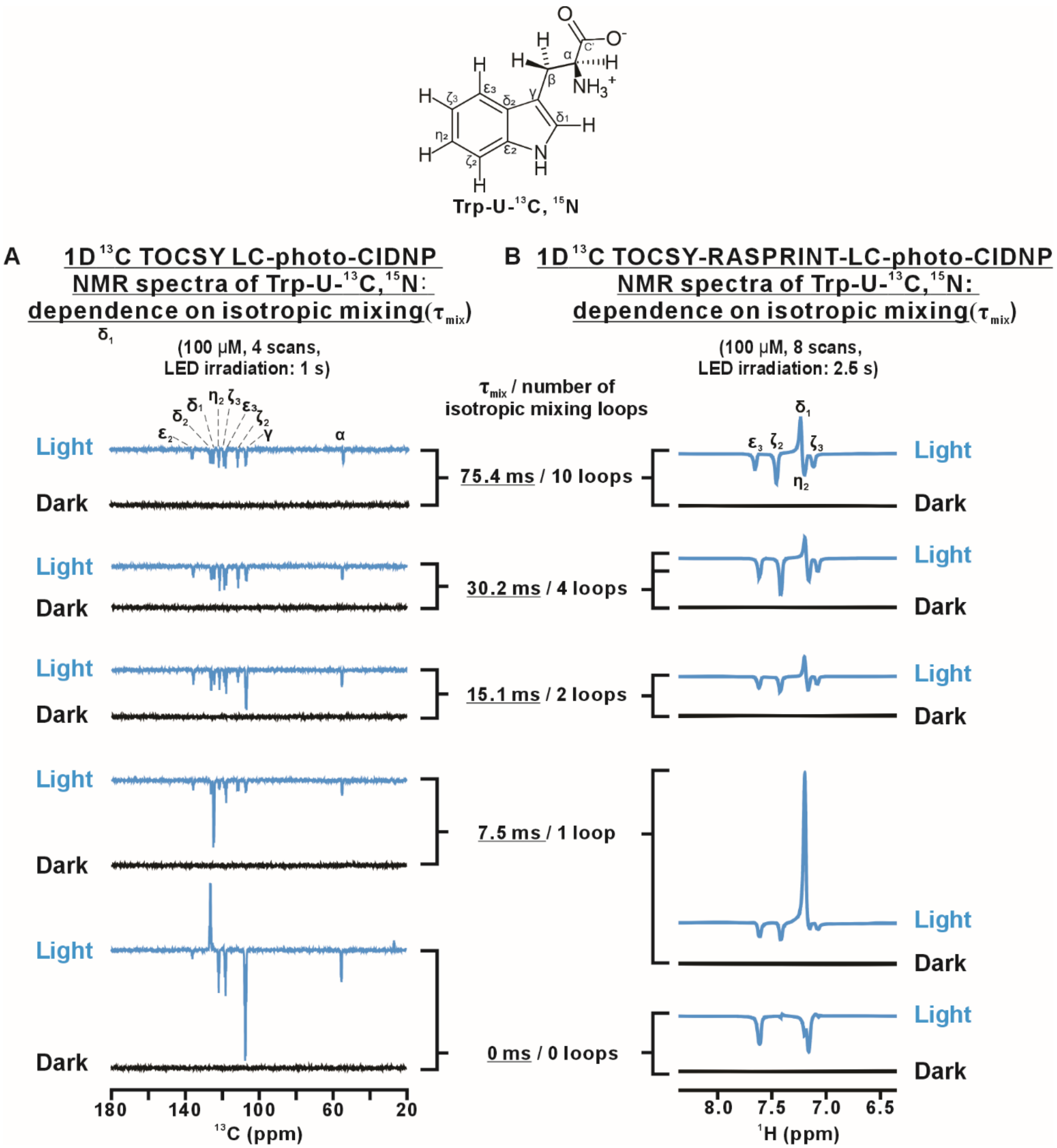


**Figure S1. Optimization of isotropic mixing time for $^{13}$C TOCSY-mediated LC-photo-CIDNP.** *(A)* $^{13}$C TOCSY LC-photo-CIDNP spectra of Trp-U-$^{13}$C, $^{15}$N acquired with increasing isotropic mixing times (rf field strength = 6,250 Hz). *(B)* $^{1}$H-detected $^{13}$C

TOCSY-RASPRINT LC-photo-CIDNP spectra acquired with different isotropic mixing times (rf field strength = 6,250 Hz). A short isotropic mixing period ($\tau_{mix}$ = 7.5 ms) provides the highest sensitivity by efficiently transferring $^{13}C$ hyperpolarization while preserving the initial polarization. Longer mixing times progressively redistribute the polarization over the entire $^{13}C$ spin system, reducing the intensity of the directly hyperpolarized resonances. The $^{1}H$-detected TOCSY-RASPRINT spectra display a similar trend. All Trp-U-$^{13}C$- $^{15}N$ samples (100 μM) were in 10 mM potassium phosphate (pH ~7.2) and included 25 μM fluorescein, oxygen-scavenging enzymes and vitamin C. See Materials and Methods for details.

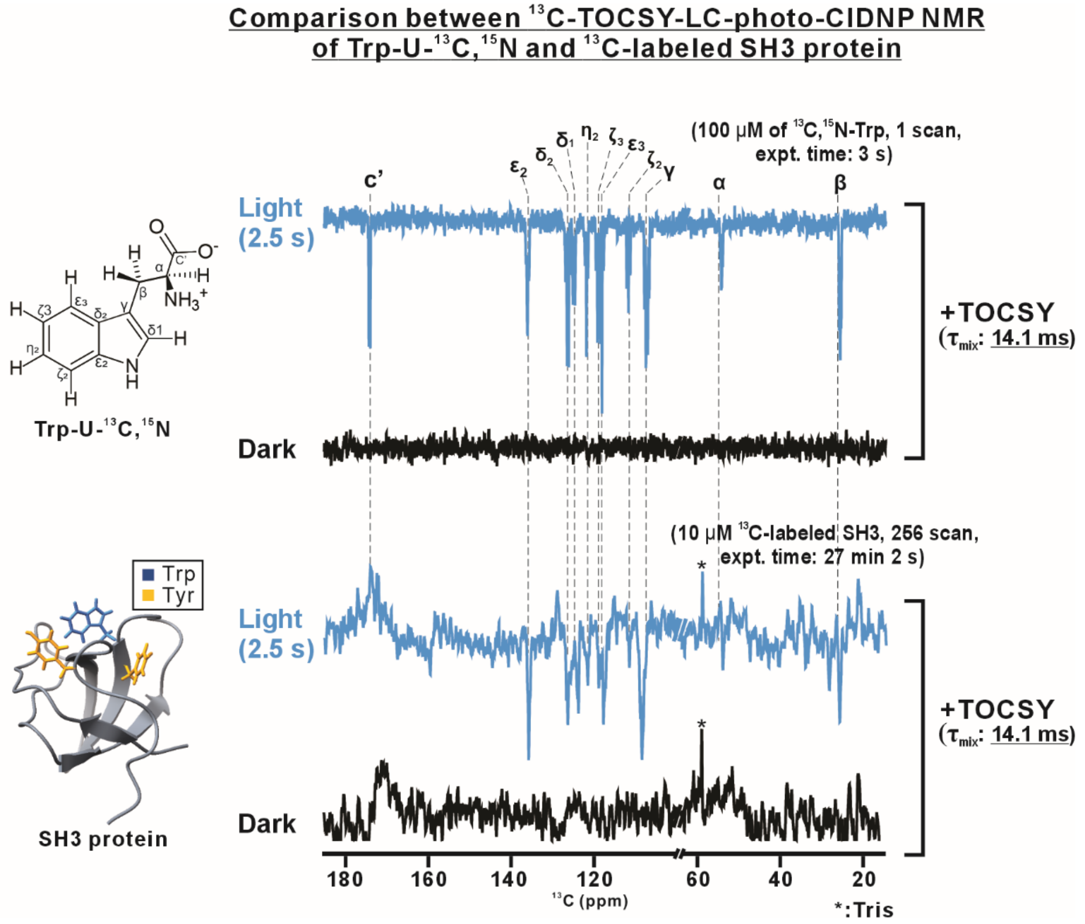


**Figure S2. Spectral comparisons between Trp-U-$^{13}$C,$^{15}$N and $^{13}$C-labeled SH3 protein.** $^{13}$C TOCSY LC-photo-CIDNP spectra of Trp-U-$^{13}$C, $^{15}$N (top) and $^{13}$C-labeled SH3 protein (bottom) acquired under identical isotropic mixing conditions (rf field = 16,666 Hz, $\tau_{mix}$ = 14.1 ms). Redistribution of the initial $^{13}$C hyperpolarization of trp-U-$^{13}$C,$^{15}$N during isotropic mixing enables observation of carbonyl ($^{13}$C′), aromatic, $^{13}$C$^{\alpha}$, and $^{13}$C$^{\beta}$ resonances. A similar polarization distribution is observed for the SH3 protein. All samples were in 10 mM potassium phosphate (pH ~7.2). The 10 μM Trp-U-$^{13}$C- $^{15}$N and 10 μM SH3 samples also contained 10 μM fluorescein, oxygen-scavenging enzymes and vitamin C. See Materials and Methods for details.

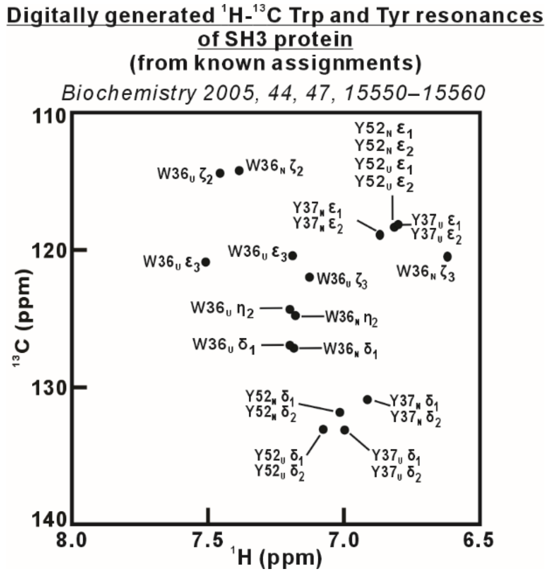


**Figure S3. Digitally generated $^{1}H$–$^{13}C$ correlation map of Trp and Tyr residues in SH3 protein derived from published NMR assignments.** *(Biochemistry 2005, 44, 15550–15560)* The map serves as a reference for identifying experimentally observed resonances and evaluating the expansion of detectable resonance coverage achieved by TOCSY-mediated LC-photo-CIDNP.
